# CRISPR/Cas9-mediated knock-out of dUTPase in mice leads to early embryonic lethality

**DOI:** 10.1101/335422

**Authors:** Hajnalka Laura Pálinkás, Gergely Rácz, Zoltán Gál, Orsolya Hoffmann, Gergely Tihanyi, Elen Gócza, László Hiripi, Beáta G. Vértessy

## Abstract

Sanitization of nucleotide pools is essential for genome maintenance. Among the enzymes significant in this mechanism, deoxyuridine 5′-triphosphate nucleotidohydrolase (dUTPase) performs cleavage of dUTP into dUMP and inorganic pyrophosphate. By this reaction the enzyme efficiently prevents uracil incorporation into DNA and provides dUMP, the substrate for *de novo* thymidylate biosynthesis. Despite its physiological significance, knock-out models of dUTPase have not yet been investigated in mammals, only in unicellular organisms, such as bacteria and yeast. Here we generate CRISPR/Cas9-mediated dUTPase knock-out in mice. We find that heterozygous *dut* +/-animals are viable while the decreased dUTPase level is clearly observable. We also show that the enzyme is essential for embryonic development. Based on the present results, early *dut* -/-embryos can still reach the blastocyst stage, however, they die shortly after implantation. Analysis of preimplantion embryos indicate perturbed growth of both inner cell mass (ICM) and trophectoderm (TE). We conclude that dUTPase is indispensable for post-implantation development in mice. The gene targeting model generated in the present study will allow further detailed studies in combination with additional gene knock-outs.

## Introduction

The maintenance of genome integrity and faithful preservation of genomic information are crucial for viability. Towards these goals, various DNA damage and repair pathways and fine-tuned regulation of well-balanced deoxynucleotide (dNTP) pool work hand-in-hand^1^. For the control of the nucleotide pools, several families of dNTP hydrolyzing enzymes are present in most organisms^2,3,4,5^. These enzymes sanitize the nucleotide pool by removing those nucleotide building blocks (dNTPs) from the polymerase action that contain erroneous bases. The dUTPase family of enzymes is responsible for the removal of dUTP from the nucleotide pool by hydrolyzing it into dUMP and inorganic pyrophosphate^6,7,8^. The importance of this enzymatic action is evident in light of the fact that most DNA polymerases cannot distinguish dUTP and dTTP and will readily incorporate the uracil-analogue if it is available in the cellular dNTP pool^9^. Through their enzymatic action that generate dUMP, dUTPases also feed into the *de novo* thymidylate biosynthesis pathway by providing dUMP as the substrate for thymidylate synthase. Two major families of dUTPases have evolved that are referred to as trimeric and dimeric dUTPases, reflecting their corresponding quaternary structure^6,10,11,12,13^.

Trimeric dUTPases are present in almost all free living organisms with the notable exception of Trypanosomes. Subunits of these enzymes contain a beta-sheeted arrangement^6^. The three subunits donate conserved sequence motifs to build the three active site of the dUTPase trimer^14,15,16^ This family of dUTPases are characteristic for Archaea, Bacteria and Eucarya. As such, mammalian species also rely on the action of trimeric dUTPases to keep a well-balanced dUTP/dTTP ratio. Herpesviruses encode an intriguing monomeric homologue of this dUTPase enzyme family^17^, where the protein sequence contains a species-specific insert to allow for constructing the usual beta-sheeted dUTPase fold in a monomeric enzyme^18,19^. Dimeric dUTPases perform the same catalytic action, however, the protein sequence and alpha-helical protein fold structure are drastically different from those observed in trimeric dUTPase family^10,11^.

Due to the highly significant enzymatic character of dUTPases, the essentiality of this enzyme family was addressed in numerous different organisms. Knock-outs have been generated in several bacteria: *Escherichia coli*^20^ and *Mycobacterium smegmatis*^7^. Based on these studies it was argued that the bacterial cells are not viable in lack of dUTPase action. However, genomic analysis of Archaea and prokaryotes identified several species that lack the dUTPase encoding gene^12^. These findings indicate that the presence of dUTPases may not be a universal requirement in prokaryotes.

The physiological role and importance of dUTPase have also been addressed in eukaryotes. In yeast, dUTPase knock-out was still viable, although this genotype led to a thymine auxotroph phenotype^21^. In *Caenorhabditis elegans,* RNA silencing studies indicated that dUTPase might be important in embryonic development^22^. Very recently, in planarians silencing of *dut* caused lethality in adult animals possibly due to genomic DNA fragmentation. Co-administration of the thymidylate synthase inhibitor 5-fluoro-uracil (5-FU) resulted in more DNA breaks and earlier planarian death^23^. In *Drosophila melanogaster,* dUTPase silencing led to early pupal lethality suggesting a specific role of dUTPase and uracil DNA metabolism in metamorphosing insects^24,25^. It has been shown that dUTPase is also essential in *Arabidopsis thaliana.* In these plants reduced dUTPase activity caused DNA damage and increased homologous recombination events. Furthermore, these plants were extremely sensitive to 5-FU^26^. In human cell lines, several laboratories published siRNA dUTPase silencing studies^27,28,29^. These all concluded in agreement that highly efficient silencing with practically no remaining dUTPase does not perturb the cellular phenotype under normal conditions^29^. Still, the dUTPase silenced cell lines showed increased sensitivity towards inhibitors of *de novo* thymidylate biosynthesis. These findings corroborated the clinical significance of dUTPase inhibition in anticancer chemotherapies^30,31,32^. To our best knowledge, knock-out studies on dUTPases have not yet been published for any mammalian species.

Motivated by the lack of knowledge in the field, we initiated dUTPase knock-out experiments in mice. Here we report successful generation of dUTPase knock-out mice using CRISPR/Cas9-mediated genome editing. We find that dUTPase knock-out leads to early embryonic lethality. No homozygous knock-out offspring could be observed, however, homozygous knock-out blastocysts are still viable and can be cultured *in vitro,* suggesting that lethality of the dUTPase knock-out sets in around or shortly after implantation.

## Results and discussion

### Generation of the dUTPase knock-out model

Fig. 1 shows the outline of the CRISPR/Cas9-mediated knock-out experiment. The respective single guide RNA was designed in such a way to disrupt both nuclear and mitochondrial isoforms of dUTPase (Fig. 1a). Efficiency of the CRISPR-mediated events were first assessed by a surveyor assay in mouse embryonic fibroblast (MEF) cells (Fig. 1b). We found evidence of CRISPR/Cas9-induced cleaved products and based on this result we started mouse zygote microinjection. Fig. 1c shows the schematics of gene targeting mice generation. In the microinjection experiments, 107 embryos were flushed and microinjected. 76 embryos were transferred to 5 foster mothers. 2 foster mothers had in sum 15 offspring. 3 out of the 15 offspring individuals were targeted (based on T7 assay), 2 were sequenced (these are numbered as #2 and #4). One of these resulted in six bp deletion and one base substitution (D6, M1), the other one resulted in 47 bp deletion (D47) leading to frameshift and early stop codons (Fig. 1d). Strains were generated from these two sequenced individuals. Homozygous and heterozygous D6, M1, as well as heterozygous D47 animals showed no gross abnormality and were fertile. The further experiments presented in this study were conducted on mouse line D47.

**Figure 1.**
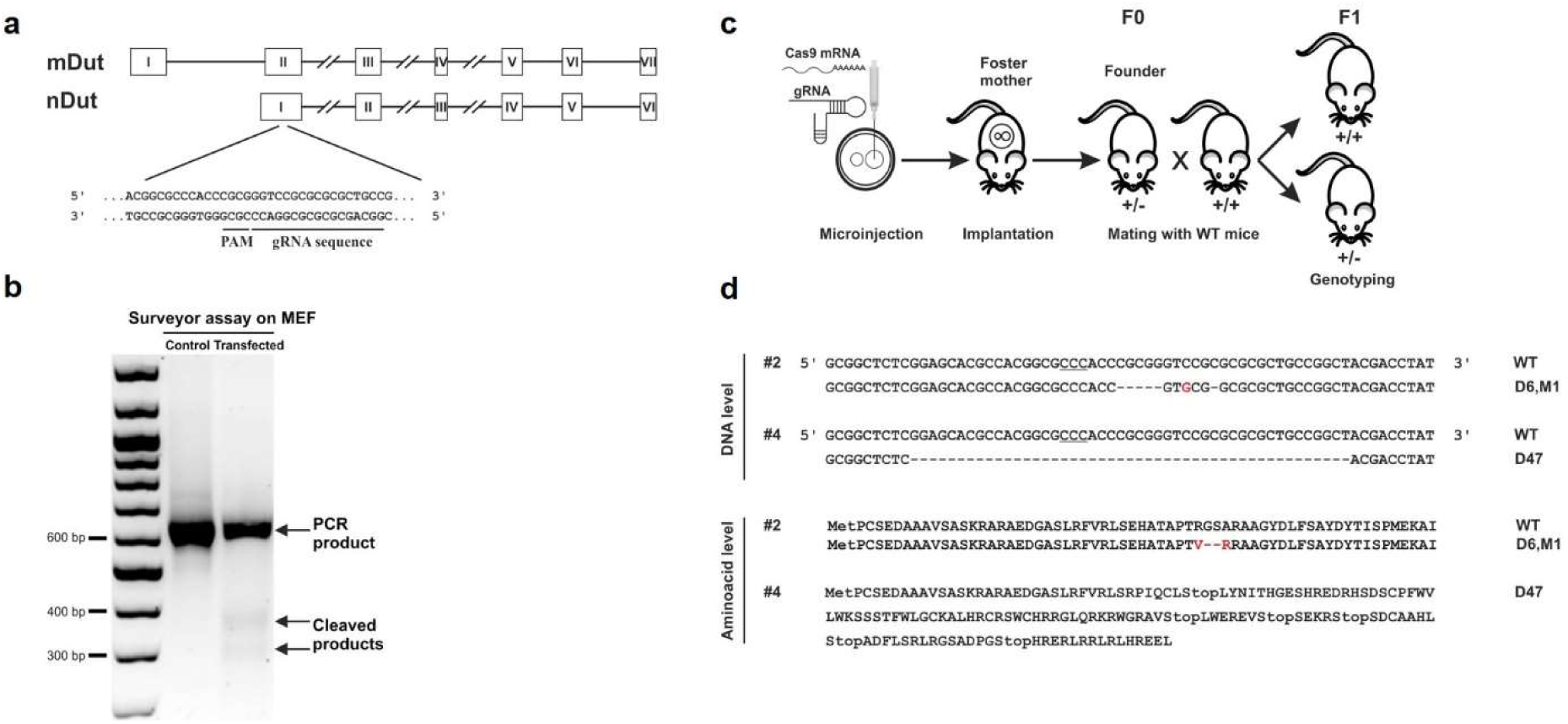
Generation and assessment of CRISPR knock-out mice. **a** Schematic diagram of the *dut* gene encoding the nuclear and mitochondrial isoforms of dUTPase. Exons are indicated with Roman numerals in rectangles, introns are depicted as lines (for longer introns lines are broken). gRNA target site and protospacer-adjacent motif (PAM) sequence in the first common exon of the two isoforms is underlined. **b** Surveyor assay performed on mouse embryonic fibroblast cells used for the detection of indel events induced by transfection with CRISPR guide RNA and Cas9 mRNA. The two lower fragments indicate cleavage of the DNA due to CRISPR events. These are lacking in the control while are visible in the transfected sample. **c** Schematic diagram showing the generation of CRISPR knock-out mice. Fertilized oocytes microinjected with CRISPR guide RNA and Cas9 mRNA were implanted into foster mothers. The resulting founders (F0) were crossbred with wild type mice to generate wild type and heterozygous offsprings (F1). **d** DNA and predicted protein sequence of the two founder mice showing CRISPR events. Mouse #2 showed deletion of 6 nucleotides and a C to G mutation (D6, M1) resulting in the deletion of two amino acids and change of another two. In mouse #4, 47 nucleotides were deleted (D47) which resulted in a frameshift mutation leading to early stop codons.

Specificity of this gRNA sequence was checked by analyzing potential off-target sites in F0 animals (Table 1). No CRISPR-induced event could be observed at these sequenced sites.

**Table 1.** Potential off-target sites predicted by CCTop -CRISPR/Cas9 target online predictor software. The Table presents the chromosomal location, the number of mismatches (MM), the target and the adjacent PAM sequence. The first row depicts the target site of the designed guide RNA, further rows list the top 10 candidates for off-target sites. Mismatches are indicated in red. Brackets include core sequences^33^.

| Name | Coordinates | MM | Target sequence | PAM |
| --- | --- | --- | --- | --- |
| Dut | chr2:125247853-125247874 | 0 | CGCGCGCGGAC[CCGCGGGT] | GGG |
| Off-1 | chr13:88680616-88680637 | 4 | TGAGTGGGAC[CCGCGGGT] | TGG |
| Off-2 | chr4:120746882-120746903 | 4 | CTGGAGCGGCC[CCGCGGGT] | GGG |
| Off-3 | chr15:28025084-28025105 | 4 | CGGCCACGGCC[CCGCGGGT] | AGG |
| Off-4 | chr11:23306808-23306829 | 4 | GGGGCGGGAG[CCGCGGGT] | GGG |
| Off-5 | chr2:118372992-118373013 | 4 | CGCTGGTGGCC[CCGCGGGT] | TGG |
| Off-6 | chr5:64803646-64803667 | 4 | ACCGCACGGAC[GCGCGGGT] | GGG |
| Off-7 | chr18:85179644-85179665 | 3 | CGTGCGCGCAC[GCGCGGGT] | GGG |
| Off-8 | chr9:77319229-77319250 | 4 | CGCGCTTACAC[CCGCGGGT] | GGG |
| Off-9 | chr2:104319532-104319553 | 3 | CGCGTGCGCAC[ACGCGGGT] | AGG |
| Off-10 | chr19:36918725-36918746 | 4 | CTCGCTGGGAC[GCGCGGGT] | AGG |

### Homozygote knock-out of dUTPase leads to embryonic lethality after the blastocyst stage

To assess the genotype after the CRISPR experiment we designed appropriate primers for the genotyping PCR reactions. These resulted in different length products from the wild type as compared to the knock-out allele in mouse line D47 (as visualized on Fig. 2a). All three genotypes (+/+, +/-, -/-) resulted in live 4.5 day-old embryos (cf. Fig. 2b).

**Figure 2.**
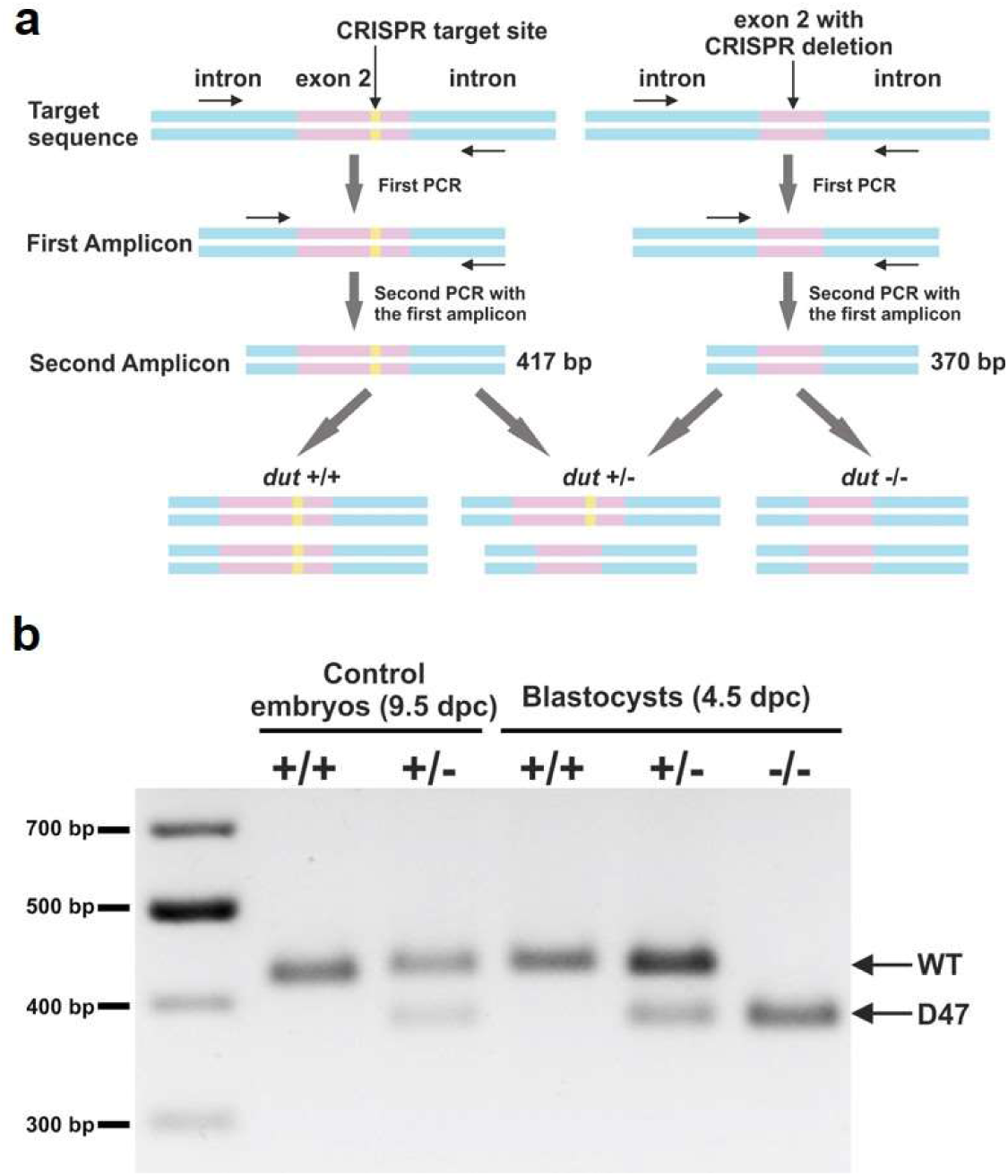
Genotyping of blastocysts. **a** Schematic representation of the used semi-nested design of genotyping. Introns are shown in blue, exons are shown in pink, and the CRISPR target site is shown in yellow. DNA isolated from blastocysts was subjected to PCR with primers (shown as arrows) adjacent to the CRISPR target site. The resulting amplicon was used in a second round of PCR with the same reverse and a nested inner forward primer to generate 417 bp length product for the WT allele and 370 bp product for D47 allele. **b** Representative image of amplicons from semi-nested PCR visualized on agarose gel. The upper and lower band corresponds to wild type and D47 allele, respectively.

Table 2 shows the results of the genotype analysis in the animals investigated in this study. The data clearly shows that *dut*-/- knock-out organisms can only be isolated in early embryonic development at the blastocyst stage.

**Table 2.** Genotype analysis of offsprings from *dut* +/-intercrosses at different developmental stages.

| DNA source | Genotype |  |  | Resorbed |
| --- | --- | --- | --- | --- |
|  | +/+ | +/- | -/- |  |
| Postnatal | 21 | 42 | 0 | NA<sup>a</sup> |
| 10.5 dpc | 3 | 5 | 0 | 3 |
| 9.5 dpc | 5 | 5 | 0 | 0 |
| 8.5 dpc | 10 | 5 | 0 | 5 |
| 3.5 dpc | 11 | 13 | 7 | NA |

<sup>a</sup> NA, not applicable

### Embryonic development in the dUTPase knock-out as compared to the heterozygous and wild type animals

dUTPase knock-out 3.5-day-old embryos from *dut* +/-intercrosses developed into normal blastocysts. After hatching, these were attached to gelatin coated surface (cf. Fig. 3). Their development were checked one day later (i.e. at 4.5 dpc (days post coitum)). It is shown in these pictures that the *dut* -/-ICM is smaller, while we do not observe obvious changes in the heterozygotes as compared to the wild type (cf. Fig. 3b). Quantitative analysis of these findings was also initiated (Supplementary Fig. S2). Further development on the attached embryos revealed that at 7.5 days, the ICM clump size is smaller in the *dut* -/-, while are rather comparable in the heterozygote and the wild type. Also, the trophoblast ectoderm was reduced in the *dut* -/-(but not in the heterozygote) (Fig. 3c).

**Figure 3.**
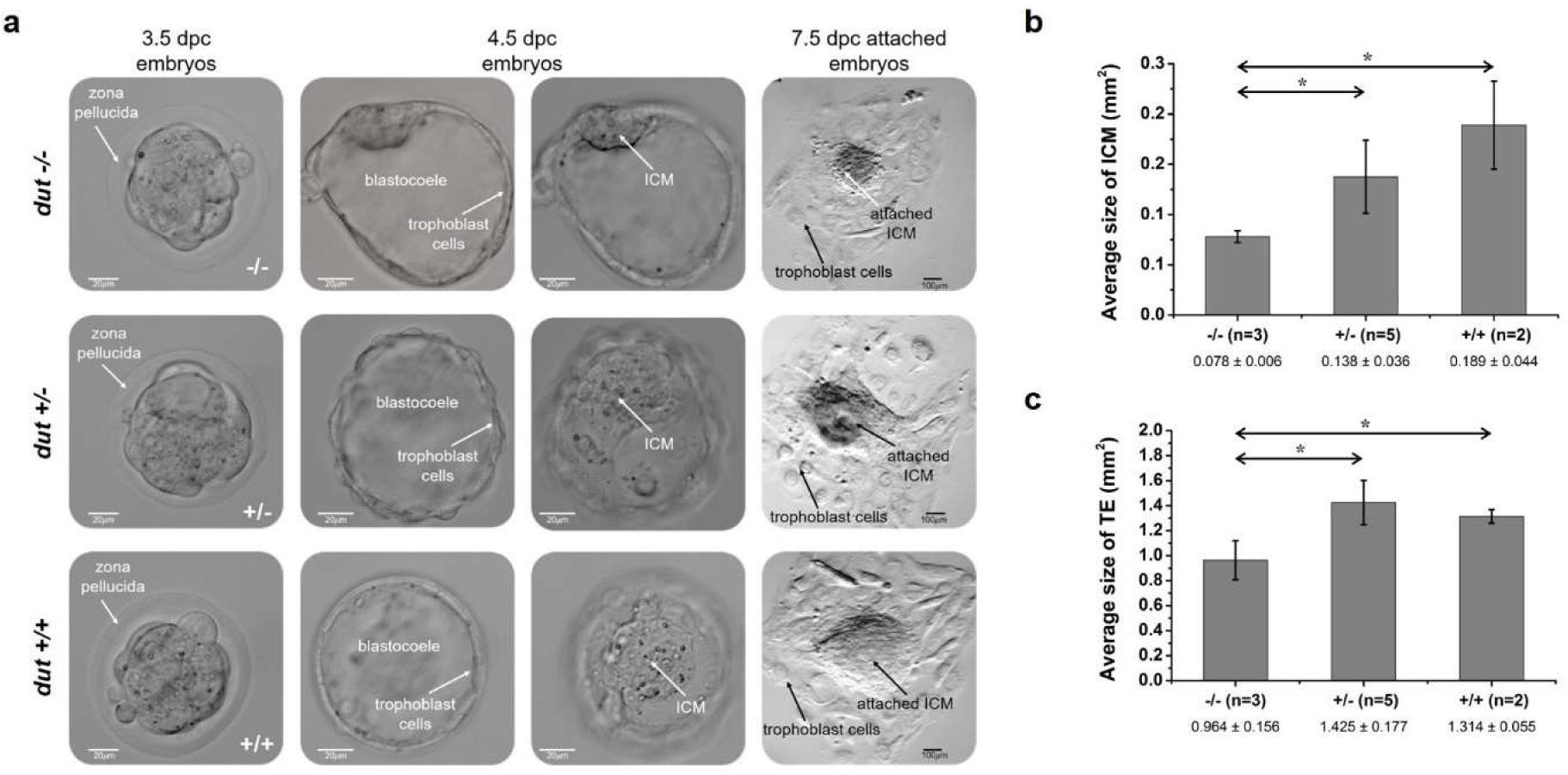
Outgrowth assay of preimplantation embryos obtained by intercrossing D47 heterozygous mice. **a** panel shows the phase contrast images of D47 homozygous (-/-), heterozygous (+/-) and wild type (+/+) blastocysts in *in vitro* culture. The first column shows embryos at 3.5 dpc after flushing from oviducts. White arrows indicate the zona pellucida surrounding the embryos. The second and third columns show the attached embryos, one day later focusing on the trophoblast cells or the inner cell mass (ICM) in the blastocoele. Scale bar, 20 μm. The last column presents outgrowths after 4 days in culture. Scale bar, 100 μm. Average size of ICM (**b**) and trophectoderm (TE) (**c**) for blastocysts of indicated genotypes. Error bars indicate standard deviation. n = 3 for (-/-), n = 5 for (+/-), and n =2 for (+/+). *, p < 0.05.

Embryos isolated from cesarean section at 8.5 dpc and 9.5 dpc are shown for wild type and heterozygotes (Fig. 4). We also observed resorbed embryos (Fig. 4a) where the genotype could not be evaluated unequivocally since the mother’s decidual tissue could not be separated from the resorptions. No further stage *dut* -/- embryos were found.

**Figure 4.**
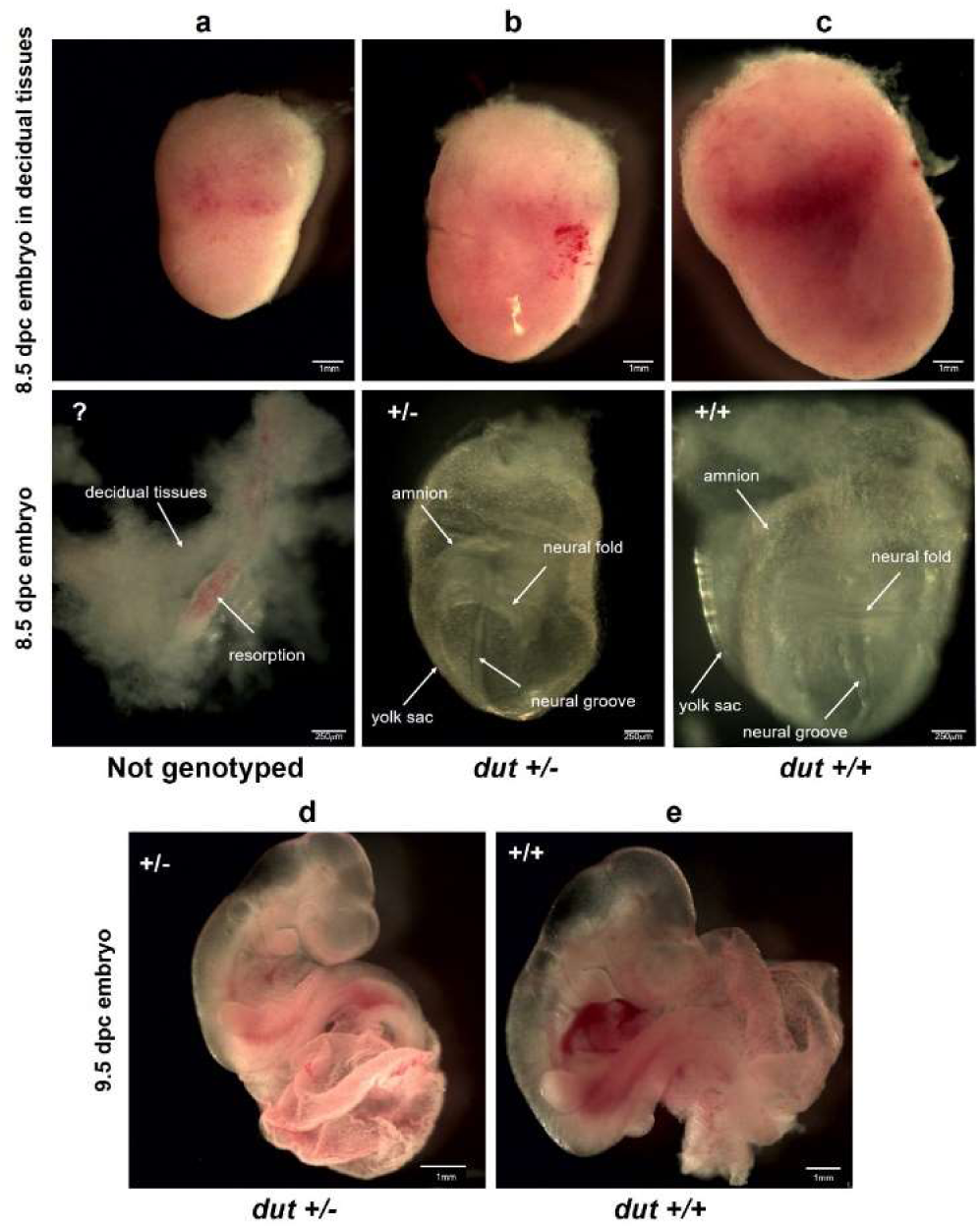
Images of embryos at 8.5 and 9.5 dpc obtained by crossing D47 heterozygous mice. **a** panel shows a resorbed embryo at 8.5 dpc covered by decidual tissues. The resorbed embryo could not be genotyped (indicated with _“_?^”^). **b** and **c** show heterozygous (+/-) and wild type (+/+) embryos at 8.5 dpc, respectively. Upper panels show embryos in intact decidual tissues. Scale bar, 1 mm. Lower panels present the embryos dissected from decidual tissues. Arrows represent the embryonic regions (neural fold, neural groove) and extra-embryonic tissues (amnion, yolk sac) of embryos. Scale bar, 250 μm. **d** Heterozygous and e wild type embryos at 9.5 dpc. Scale bar, 1 mm.

To analyze the protein presence in the wild type and heterozygous mouse, we performed Western blot analysis (Fig. 5). The dUTPase protein level was clearly reduced in the heterozygote as compared to the wild type as discussed above. *Dut* -/- homozygous embryo could not be found at this embryonic stage.

**Figure 5.**
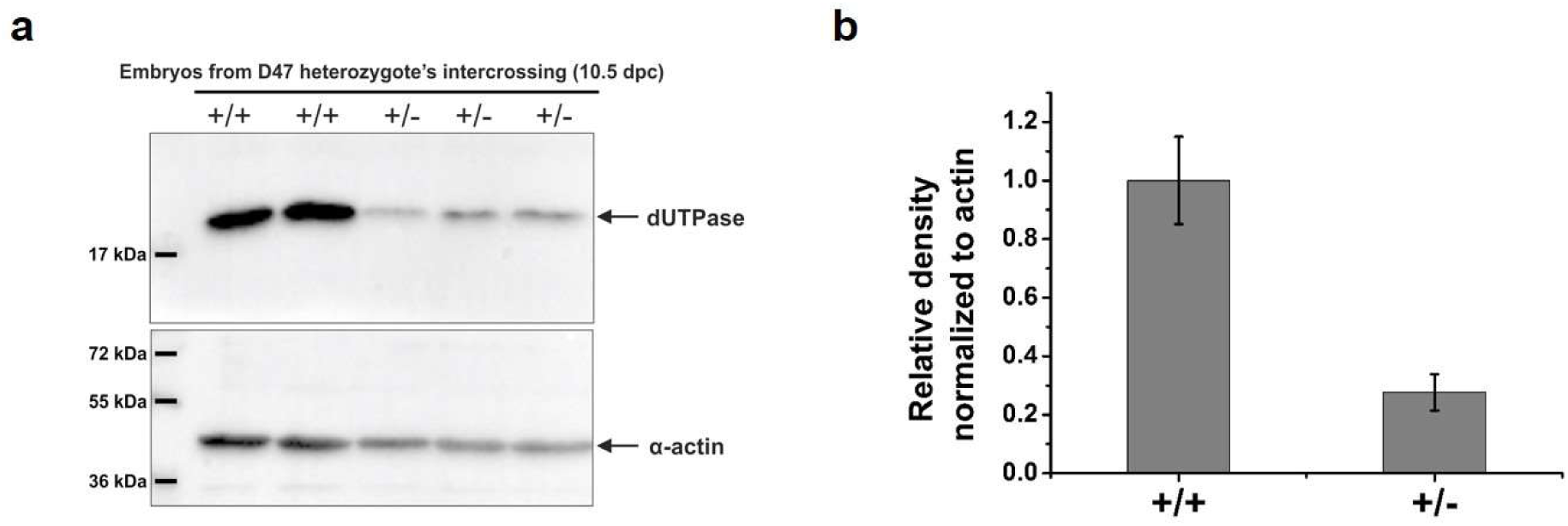
Protein level of dUTPase in embryos at 10.5 dpc from intercrossing of D47 heterozygous mice. **a** Western blot displaying dUTPase protein level in wild type and heterozygous mice. α-actin serves as loading control. **b** Densitometric data for dUTPase levels from Western blot normalized for α-actin and the mean of wild types. Error bars indicate standard deviation. n = 2 for (+/+) and n = 3 for (+/-).

### Concluding remarks

The schematic pattern of embryonic development in the mice is illustrated on Figure 6. In this scheme, the timeline of the degradation of the maternal mRNA and protein and the parallel activation of the zygotic genome is also indicated^34,35^. Our gene targeting results showed that the dUTPase knock-out allowed the first several duplication cycles to proceed and viable blastocysts could be isolated. Also, isolated *dut* -/-blastocysts could grow further in *in vitro* cultures. However, further development following implantation was prevented in the homozygous knock-out.

**Figure 6.**
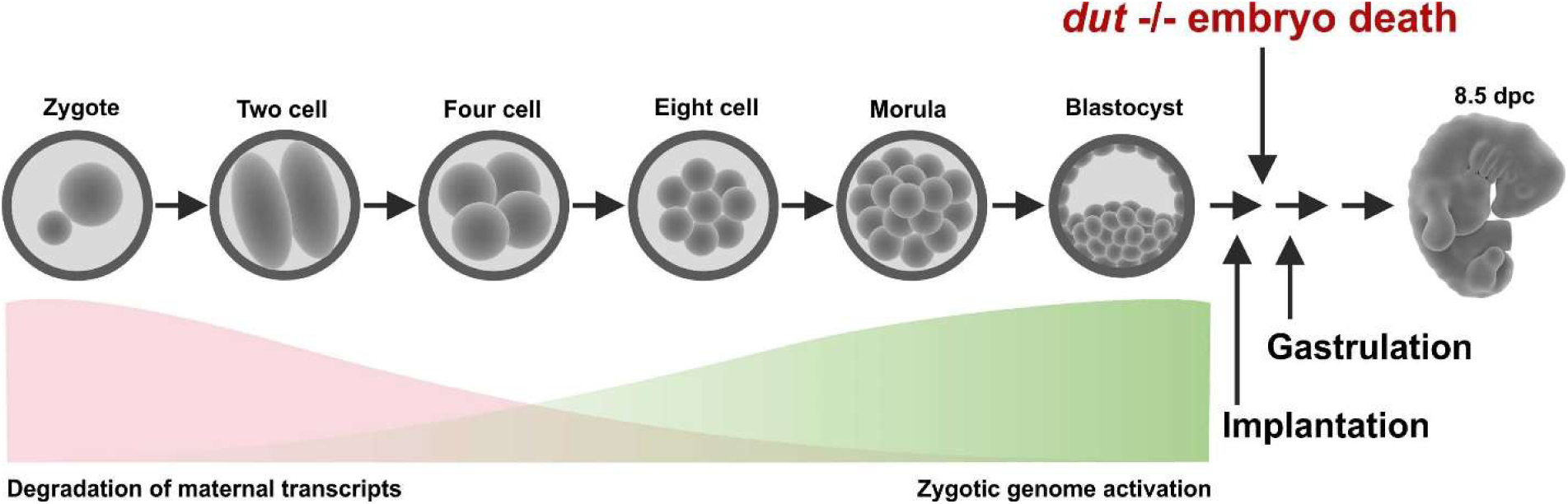
Schematic diagram of embryonic development in mice. The red and green areas show the timeline of simultaneous degradation of maternal transcripts and activation of zygotic transcription. Arrows illustrate that homozygous knock-out embryos die shortly after implantation.

We conclude that mitotic events may proceed in lack of dUTPase in mice, however, the enzyme is indispensable for later differentiation stages. Our model will be useful in later detailed studies to outline the molecular events leading to the observed early embryonic lethal phenotype. In addition, combined multiple knock-outs of *dut* and other relevant genes are also expected to provide important insights into developmental processes.

## Methods

### CRISPR constructs

The T7 single guide RNA (sgRNA) harboring the PAM sequence and the Cas9 mRNA were obtained from Sigma-Aldrich. According to our request, the designed sgRNA targets the first common exon of dUTPase isoforms (exon 2) in the mouse genome (cf. Fig. 1a).

### CRISPR/Cas9 efficiency test in MEF cells

Target sgRNA and Cas9 nuclease mRNA were transfected into mouse embryonic fibroblast (MEF) cells by Lipofectamine™ 3000 transfection reagent (ThermoFisher Scientific). According to manufacturer’s recommendation, 2.5 μg Cas9 mRNA and 250 ng target sgRNA were added to the sub-confluent cultures of cells grown in 6-well plates. After 24 hours post-transfection, cells were maintained in a fresh medium for 24 hours, and then the genomic DNA was extracted with MasterPure™ DNA Purification Kit (Epicentre, Madison, WI, USA). After DNA amplification with Cel-1-F and Cel-1-R primers, Cel 1 cleavage assay was performed using the Transgenomic® SURVEYOR® Mutation Detection Kit according to manufacturer’s instructions. The primers used in this study were synthesized by Sigma-Aldrich, and are listed in Supplementary Table S1.

### Animals

dUTPase wild type and heterozygous mice used in the experiments were produced and maintained in the Animal Care Facility at the NAIK Agricultural Biotechnology Center (FVB/N background, Envigo UK). Animals were housed in groups of 2–5 with free access to food and water. Animals were kept under standard light-dark cycle (06.00–18.00 h) at 22°C. This study was carried out in strict accordance with the recommendations and rules in the Hungarian Code of Practice for the Care and Use of Animals for Scientific Purposes. The protocol was approved by the Animal Care and Ethics Committee of the NAIC-Agricultural Biotechnology Institute and the Pest County’s governmental office (permission number: PEI/001/329-4/2013). The method used for euthanasia: cervical dislocation. All efforts were made to minimize suffering.

### Micromanipulation and detection of gene targeting

Microinjection was performed as described^36^. Briefly, mouse zygotes were collected from superovulated FVB/N (Envigo, UK) females mated with FVB/N males. Pronuclei were injected using a manual injector with continuous flow. Following visualization of pronuclear swelling, the needle was pulled out through the cytoplasm, injecting a small amount of additional RNA delivery to the cytoplasm. The microinjection mix contained a sgRNA (Sigma, USA) diluted in 10 mM Tris, 0.2 mM EDTA (pH = 7.4) in a final concentration of 10 ng/μl and Cas9 mRNA (Trilink, USA) in 10 mM Tris, 0.2 mM EDTA (pH = 7.4) in a final concentration of 150 ng/μl. Injected zygotes were transferred to pseudopregnant CD-1 females (Envigo, UK). All animals born from embryo transfer were genotyped by PCR and T7 assay.

### Cloning and sequencing

Genotyping of the heterozygous founder animals was carried out by amplifying the CRISPR target sites from genomic DNA using primers pBS-F and pBS-R (listed in Supplementary Table S1), and the fragments were cloned into *SalI/EcoRI* sites of vector pBluescript SK (+) (Stratagene). 20 of individual bacterial colonies were purified with NucleoSpin^®^ Plasmid DNA Purification Kit (MACHEREY-NAGEL GmbH & Co. KG) according to manufacturer’s instructions, then DNA samples were subjected for sequencing. Based on the sequencing results, two animals (founder #2 and #4) showed CRISPR events, whose offsprings were termed as D6 and D47, respectively. All DNA samples in this study were verified by sequencing by Microsynth Seqlab GmbH.

### Off-target analysis

Off-target effects of CRISPR/Cas9 nucleases were evaluated via the online predictor CCTop -CRISPR/Cas9 target (https://crispr.cos.uni-heidelberg.de/)^33^. Ten candidate loci for the target site with high potential cleavage in mouse genome were chosen. The selected potential off-target sites were PCR-amplified using genomic DNA from founder animal #4 and evaluated by DNA sequencing. Eight of ten candidates could be analyzed, whereas two of them could not be evaluated due to aspecific PCR product from repetitive elements. The information on the off-target loci identified by the online program are shown on Fig. 1e and sequencing primer pairs used are listed in Supplementary Table S1.

### Genotyping

The genotypes of mice were determined by PCR of total genomic DNA extracted from mouse tails. Genotyping of embryos was also performed by PCR either by isolating DNA from full embryos or from outgrowth assays. All isolated samples were dissolved in DNA lysis buffer (0.1 M TRIS-HCl pH = 7.4; 0.2 M NaCl; 5 mM EDTA; 0.2% SDS) and DNA was extracted with phenol-chloroform. MyTaq polymerase (Bioline) was activated at 95°C for 5 min, and PCR was performed for 30 cycles at 95°C for 30 s, 64°C for 30 s, and 72°C for 30 s, with a final extension at 72°C for 10 min using primers Dut-gen-F and Dut-gen-R (Supplementary Table S1). Genotyping from blastocysts was performed by semi-nested PCR using 1 μl template from 30x diluted primary PCR product with primers Dut-nest-F and Dut-gen-R (Supplementary Table S1) under the same reaction conditions. DNA fragments were visualized by 1% agarose gel electrophoresis.

### Analysis of dissected embryos

For the analysis presented in this manuscript, D47 heterozygous males were mated with D47 heterozygous females, and embryos at various stages (3.5-9.5 dpc) were collected from pregnant D47 heterozygous females. Dissections were performed in ice-cold PBS. After dissection embryos were examined and photographed with LeicaM205FCA-FC fluorescent stereo microscope linked to a DFC7000-T Leica Camera. The day of plug formation was defined as embryonic day 0.5.

### Analysis of blastocysts outgrowths

Embryos were flushed out from uteri of pregnant mice at 3.5 dpc in M2 medium. Blastocysts were individually cultured on 0.1% gelatin-coated, 12-well tissue culture dishes (Eppendorf), in KO-DMEM ES cell culture medium supplemented with 1000 U/ml LIF and 20% fetal bovine serum (HyClone), in 5% CO2 at 37°C for 4 days. Outgrowth were photographed daily using LeicaM205FCA-FC fluorescent stereo microscope linked to a DFC7000-T Leica Camera. On the fourth days of culture, outgrowth were photographed, subsequently removed and genotyped by PCR as described above.

### Western blot

Embryos at 10.5 dpc were dissected immediately following euthanasia of pregnant mice, then washed with PBS and resuspended in lysis buffer (20 mM HEPES pH = 7.5; 420 mM NaCl; 1 mM EDTA; 2 mM dithiothreitol (DTT); 25% glycerol). Homogenization was assisted with vortex until the tissue was sufficiently disrupted. Samples were centrifuged at 20.000 g for 15 min at 4°C to remove insoluble fraction, then the supernatant samples were boiled with SDS buffer at 95°C for 5 min. Proteins were resolved under denaturing and reducing conditions on a 12% polyacrylamide gel and transferred to PDVF membrane (Immobilon-P, Merck Millipore, Billerica, MA, USA). Membranes were blocked with 5% non-fat dried milk in TBS-T (25 mM TRIS-HCl pH = 7.4; 140 mM NaCl; 3 mM KCl; 0.05% Tween-20) for 1 h at 4°C and were developed against dUTPase (1:2000, Sigma-Aldrich) and α-actin (1:1000, Sigma-Aldrich) for loading control. After applying horseradish peroxidase coupled secondary antibodies (Amersham Pharmacia Biotech), immunoreactive bands were visualized by enhanced chemiluminescence reagent (Millipore, Western Chemiluminescent HRP substrate) and images were captured by a BioRad ChemiDoc™ MP Imaging system. Densitometry was done using BioRad Image Lab™ 6.0.

## Acknowledgements

Supported by the National Research, Development and Innovation Office of Hungary (K109486, K119493, NVKP_16-1-2016-0020, 2017-1.3.1-VKE-2017-00002, 2017-1. 3.1-VKE-2017-00013, VEK0P-2.3.2-16-2017-00013 to BGV). This work was supported by GENNET_21 (VEK0P-2.3.2-16-2016-00012 to EG). HLP was also supported by the UNKP-17-3 New National Excellence Program of the Ministry of Human Capacities.

## Author contributions statement

HLP, EG, LH and BGV conceived the experiments, HLP, GR, ZG, OH, GT, EG and LH conducted the experiments, HLP, GR, ZG, GT, EG, LH and BGV analyzed the results. All authors contributed to manuscript writing.

## Data availability statement

All data generated or analyzed during this study are included in this published article (and its supplementary information files).

